# Structural Basis for Katanin Self-Assembly

**DOI:** 10.1101/197558

**Authors:** Stanley Nithiananatham, Francis J. McNally, Jawdat Al-Bassam

## Abstract

The reorganization of microtubules in mitosis, meiosis and development requires the microtubule-severing activity of katanin. Katanin is composed of a AAA ATPase subunit and a regulatory subunit. Microtubule severing requires ATP hydrolysis by katanin’s conserved AAA ATPase domains. Whereas other AAA ATPases form stable hexamers, we show that wild-type katanin only forms heterodimers and heterotetramers. Heterododecamers were only observed for an ATP hydrolysis deficient mutant in the presence of ATP, suggesting an auto-inhibition mechanism that prevents oligomerization. X-ray structures of katanin’s AAA ATPase in monomeric nucleotide-free and pseudo-oligomeric ADP-bound states reveal conformational changes in AAA subdomains and N and C-terminal expansion segments that explain this auto-inhibition of assembly. These data lead to a model in which self-inhibited heterodimers bind to a microtubule, then transition into an assembly-competent conformation upon ATP binding. Microtubule-bound heterododecamers then promote tubulin extraction from the microtubule prior to oligomer dissociation.

## INTRODUCTION

Microtubules (MTs) are dynamic cytoskeleton polymers that are essential force generators that organize the cytoplasm during cell division, development and morphogenesis. The stability of the MT polymer is mediated by longitudinal and lateral interfaces between αβ-tubulins polymerized in the MT lattice. Diverse classes of MT regulators promote the polymerization and depolymerization of dynamic MTs by binding αβ-tubulins at their ends. However, in contrast to these regulators, MT severing enzymes destabilize MTs by binding along MT lattice sites and generate several new MTs. The MT severing proteins include the closely related AAA ATPases katanin, spastin and fidgetin, which are conserved across protozoa, plants and metazoans (Roll-Mecak and McNally, 2010). MT severing enzymes carry out essential MT regulatory functions in many cellular settings in which MT functions are involved. During mitosis and meiosis, they activate MT disassembly (Jiang et al., 2017; McNally et al., 2006). During neuronal development, they are essential to release new MTs after nucleation (Ahmad et al., 1999), and during cell motility they regulate MT formation in cilia or flagella (Sharma et al., 2007). Defects in human katanin and spastin lead to neurological disorders such as microlissencephaly (Hu et al., 2014; Mishra-Gorur et al., 2014) or hereditary spastic paraplegia (Hazan et al., 1999), respectively.

Katanin was first purified from sea urchin eggs and is composed of a catalytic subunit termed p60 and a regulatory subunit termed p80 (McNally and Vale, 1993). p60 katanin is composed of an N-terminal microtubule-interacting and trafficking (MIT) domain (Iwaya et al., 2010), followed by a 50-70-residue linker and a highly conserved Cterminal AAA ATPase domain. p80 katanin exists in multiple forms, which either include or exclude a large N-terminal β-propeller or WD-40 domain, followed by a core conserved 280-residue helical bundle region termed Conserved p80 (con80) (Jiang et al., 2017) (Figure 1A). Both p60 (MIT and AAA domains) and p80 (con80 with or without WD-40) domains are essential for MT severing function. p60 and p80 katanin form a complex through con80-MIT domains, the structure of which reveals a helical assembly which is responsible for binding MTs and recruiting MT regulatory factors to sites of MT lattice deformation (Jiang et al., 2017). p60 and p80 katanin co-purify from many cell types (McNally and Thomas, 1998; McNally and Vale, 1993), and p60 and p80 mutants have identical phenotypes in many organisms (Sharma et al., 2007; Srayko et al., 2000), indicating a complex of p60 and p80 is the relevant physiological complex.

**Figure 1:**
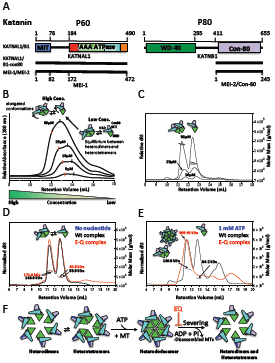
Domain structures and biochemical characterization of katanins. (A) Schematic representation of *Homo sapiens* Katanin, a heterodimer of P60 and P80 proteins (KATNAL1/KATNB1) that consist of MIT (blue), AAA (red, N-Helix; light green, NBD; cyan, HBD and orange, C-Helix), WD-40 (green) and con-80 (light purple) domains, respectively. The KATNAL1/B1-con80 and *Caenohabditis elegans* MEI-1/MEI-2 constructs are shown below. (B) SEC based analysis of wild-type KATNAL1/B1-con80 at concentrations ranging from 5 to 50 μM in the presence of 1 mM ATP using size-exclusion chromatography (Superose-6 column). Note that the complex migrates more slowly at lower concentrations, suggesting that KATNAL1/B1-con80 is in equilibrium. (C) SEC-MALS of wild-type KATNAL1/B1-con80 at the same concentrations in the presence of 1 mM ATP using superdex-200 column show heterodimers and heterotetramers, revealing that the complex is in an equilibrium. (D) SEC-MALS of wild-type (black) and E308Q mutant (red) of KATNAL1/B1-con80 reveal the complexes show in a heterodimer-heterotetramer equilibrium in the absence of 1 mM ATP. (E) SEC-MALS curves reveal the KATNAL1 (E308Q)/B1-con80 (red) forms heterododecameric assembly in the presence of 1 mM ATP, but only hetrotetrameric assembly with wild-type KATNAL1/B1-con80 (black). (F) Schematic reaction scheme Katanin p60-p80 oligomerization and in role of ATP binding and hydrolysis in this process.

Most AAA ATPases are thought to act as hexameric ring complexes. For example, Nethylmaleimide sensitive fusion (NSF) is greater than 90% hexamer in solution throughout a concentration range of 0.2 to 10 μM in ATP (Fleming et al., 1998). X-ray or cryo-EM structures of several other wild-type AAA ATPases have revealed open or closed hexameric rings (DeLaBarre and Brunger, 2005; Fletcher et al., 2003; Suno et al., 2006). In contrast, members of one sub-family of AAA ATPases have been reported to be predominantly monomeric or dimeric in solution. This sub-family includes the MT-severing proteins katanin, spastin and fidgetin, as well as Vps4 and MSP1, which disassemble non-tubulin substrates (Babst et al., 1998; Hartman and Vale, 1999; Roll-Mecak and Vale, 2008; Wohlever et al., 2017). Each of these ATPases has been reported to form a stable hexamer at low concentration only when ATP hydrolysis is blocked by mutation of a conserved E in the Walker B motif to Q (Babst et al., 1998; Hartman and Vale, 1999; Roll-Mecak and Vale, 2008; Wohlever et al., 2017) and the wild-type version of each has most often been shown to be monomeric in solution (Roll-Mecak and Vale, 2008; Scott et al., 2005; Wohlever et al., 2017).

Here, we reconstitute complexes of full-length p60 with core con80 domains for *C. elegans* MEI-1/MEI-2 and human KATNAL1/KATNB1. We show that p60-p80 is a heterodimer or heterotetramer with a poor capacity for oligomerization. Crystal structures for katanin’s AAA-ATPase reveal a substantial conformational transition for the monomeric AAA-ATPase subunit to form oligomeric assemblies. These data reveal a unique self-regulated conformation in the katanin AAA-ATPase domain that regulates its oligomerization and suggest a model for MT severing involving regulated monomer assembly, followed by ring assembly in the presence of ATP and MTs.

## RESULTS

### Catalytically active p60/p80 katanin is predominantly composed of heterodimers or heterotetramers in solution

To examine the oligomerization properties of p60/p80 katanin, we first purified complexes of full-length *C. elegans* MEI-1 with full-length MEI-2 (termed MEI-1/MEI-2), and full length human KATNAL1 with the con80 domain (residues 411-655) of human KATNB1 (termed KATNAL1/B1-con80) (Figure 1A, Figure S1A,B). KATNAL1/B1-con80 complexes were studied using a Superose 6 size exclusion (SEC) column at katanin concentrations ranging from 5 to 50 μM in the presence of 1 mM ATP (Figure 1B). Two overlapping peaks eluted earlier at higher concentrations, indicating a concentration-dependent increase in Stoke’s radius. To test whether this increase in Stoke’s radius was due to an increase in mass due to oligomerization of heterodimers, 10-50 μM KATNAL1/B1-con80 complexes were analyzed by SEC-MALS (multi angle light scattering) in the presence of 1 mM ATP using a Superdex 200 column (Figure 1C; Table 1). The two peaks of katanin complexes were more clearly resolved using the Superdex-200 SEC column. SEC-MALS measurements indicated a mass for the faster eluting complex that was most consistent with a dimer of heterodimers (heterotetramer) and a mass for the slower eluting complex that was most consistent with a heterodimer (Figure 1D). Strikingly, these masses did not change in the range of 10 - 50 μM katanin (Table 1) even though the Stoke’s radii increased in this concentration range (Figure 1B,C). For wild-type KATNAL1/B1-con80, these masses were identical in the absence (Figure 1D) or in the presence (Figure 1E; Table 1) of ATP. In the absence of ATP, *C. elegans* MEI-1/MEI-2 eluted as a single peak with a mass most consistent with a dimer of heterodimers (heterotetramer) (Figure S1C; Table 1). Previous studies of katanin (Hartman and Vale, 1999; Zehr et al., 2017) and spastin (Roll-Mecak and Vale, 2008) indicated that an E-Q mutation in the Walker B motif stabilized a hexameric assembly only in the presence of ATP. Indeed an E308Q variant of KATNAL1/B1-con80 had a mass most consistent with a heterododecamer or a hexamer of heterodimers only in the presence of ATP (Figure 1D, E; Table 1). In the absence of ATP, KATNAL1 (E308Q)/B1-con80 eluted in two peaks with masses consistent with heterodimers and heterotetramers, similar to wild-type KATNAL1/B1-con80. Strikingly, the Stoke’s radius of a KATNAL1 (E308Q)/B1-con80 hexamer of heterodimers (heterododecamer) was nearly identical to that of a wild-type dimer of heterodimers at the same concentration (Figure 1E). Previous work estimated the *in vivo* concentration of katanin at 20-50 nM (McNally and Thomas, 1998). These results thus indicate that wild-type katanin does not form hexamers at concentrations one thousand times higher than physiological concentrations and suggest that in solution, wild-type katanin is in an autoinhibited state. Furthermore, the E308Q mutation does not affect the katanin autoinhibited state in the absence of ATP.

**Table 1:**
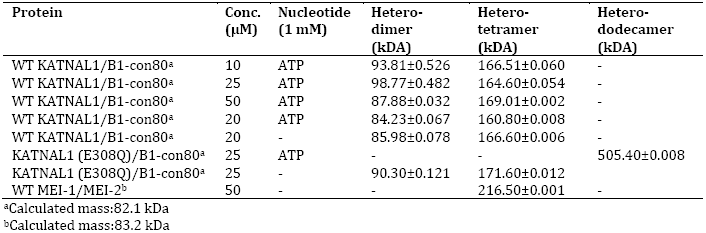
The oligomeric assembly activities of Katanin.

### Crystal structure of katanin’s AAA-ATPase in the ADP-bound state reveals a pseudo-hexameric left-handed spiral assembly

To understand the mechanism for katanin assembly, we aimed to determine the crystal structure of katanin with AMPPNP. Although we crystalized the full length *C. elegans* p60 katanin in complex with the con80 domain of p80, MEI-1/MEI-2 crystals only contained the AAA ATPase domains and bound ADP molecules. We observed degradation of MIT-Con80 domains during crystallization likely due to sensitivity to proteolysis at the AAA-MIT linker region. Crystals of *C. elegans* MEI-1/MEI-2 formed in the presence of ADP in the space group *P*6_5_ and diffracted to 3.1-Å resolution. Phase information was determined by single anomalous dispersion (SAD) using selenomethionine substituted protein (see STAR Methods). The refined 3.1-Å structure includes only part of the AAA-MIT linker region (residues 164-171) and the MEI-1 AAA-domain starting at residue 172 to residue 468 (Figure 2A,B; Table 2) in which three flexible loops are disordered (see STAR Methods). The remaining MEI-1 and MEI-2 regions are likely degraded during the crystallization process. The structure reveals an AAA-ATPase fold with a larger nucleotide binding domain (NBD) and a smaller four-helix bundle domain (HBD) (Figure 2B). The large domain is composed of a five-stranded β-sheet (β1-5) forming a parallel sheet sandwiched between nine α-helices (α2-10). The NBD forms an α/β Rossman fold that cradles an ADP molecule via Walker A and Walker B motifs (Figure 2E,F). The HBD is a helical domain composed of a central α-helix bound orthogonally by four anti-parallel helices (α11-13, α16) with its sensor II motif. Two additional short helices (α14-15) were inserted between α13 and α16. This insertion is replaced by the β domain in the Vps4 AAA ATPase (Scott et al., 2005). The MEI-1 AAA structure reveals two highly conserved katanin expansion AAA-ATPase segments, which we term the N-terminal (α1; N-Helix or N-Hlx) and C-terminal (α17; C-Helix or C-Hlx) helices, respectively (Figure 2A,B and Figure S2). The N-helix consists of single turn helix bound along one side of the NBD triangle followed by a linker and three-turn helix bound along the opposing edge of the NBD triangle (Figure 2B). The C-helix stabilizes the NBD and HBD junction, which are connected by a short loop we term the hinge. The MEI-1 AAA structure reveals the conformation of the AAA subdomains in a pseudo-oligomeric assembly state. In this structure, the protomers are arranged along a pseudo-hexameric staircase-like screw axis with one subunit in the asymmetric unit (Figure 2C). The projection of this structure parallel to this crystallographic screw axis reveals a pseudo-hexameric left-handed spiral assembly with a 14-Å translation between two adjacent subunits (Figure S3A,B). In this conformation, the NBD and ADP of one AAA subunit is bound by an HBD via the sensor II motif with 1112 Å2 buried surface per monomer that might reflect the functional contacts of a physiologically relevant oligomer (Figure 2C,E,F). The expansion segments (N-Hlx and C-Hlx) line opposite sides of this left-handed katanin spiral.

**Table 2:** Crystallographic data collection and refinement statistics.

| Data collection | KATNAL1 (E308Q) AAA | KATNAL1 (E308Q) AAA<br>gold derivative (Peak) | MEI-1 AAA<br>(SeMet peak) |
| --- | --- | --- | --- |
| Resolution range (Å) | 61.19- 2.40 (2.53- 2.40)<sup>a</sup> | 60.88-3.90 (4.11-3.90) | 85.59 – 3.10 (3.27 – 3.10)<sup>a</sup> |
| Space group | *P* 2<sub>1</sub> 2<sub>1</sub> 2<sub>1</sub> | *P* 2<sub>1</sub> 2<sub>1</sub> 2<sub>1</sub> | *P* 6<sub>5</sub> |
| Wavelength (Å) | 0.9792 | 1.0388 | 0.9791 |
| Unit cell (Å): *a*, *b*, *c* | 40.10, 61.19, 117.62 | 40.30, 60.88, 118.70 | 98.83, 98.83, 75.08 |
| Total number of observed reflections | 73593 (7317)<sup>a</sup> | 11699 (1763)<sup>a</sup> | 37571 (5592)<sup>a</sup> |
| Unique reflections | 11932 (1713)<sup>a</sup> | 2897 (414)<sup>a</sup> | 7653 (1118)<sup>a</sup> |
| Average mosaicity | 0.71 | 0.45 | 0.85 |
| Anomalous Multiplicity | - | 2.3 (2.3)<sup>a</sup> | 2.4 (2.4)<sup>a</sup> |
| Multiplicity | 6.2 (4.3)<sup>a</sup> | 4.0 (4.3)<sup>a</sup> | 4.9 (5.0)<sup>a</sup> |
| Anomalous Completeness (%) | - | 92.9 (95.6)<sup>a</sup> | 94.7 (94.6)<sup>a</sup> |
| Completeness (%) | 99.9 (99.4)<sup>a</sup> | 98.8 (99.2)<sup>a</sup> | 99.6 (100.0)<sup>a</sup> |
| $\langle I/\sigma(I) \rangle$ | 10.4 (2.1)<sup>a</sup> | 4.7 (3.0)<sup>a</sup> | 13.0 (2.4)<sup>a</sup> |
| $R_{\text{merge}}^b$ | 0.094 (0.569)<sup>a</sup> | 0.18 (0.42)<sup>a</sup> | 0.076 (0.624)<sup>a</sup> |
| **Structure refinement** |  |  |  |
| $R_{\text{work}}$ | 0.21 (0.27)<sup>a</sup> | - | 0.20 (0.24)<sup>a</sup> |
| $R_{\text{free}}$ | 0.27 (0.31)<sup>a</sup> | - | 0.27 (0.35)<sup>a</sup> |
| Molecules per asymmetric unit | 1 | - | 1 |
| Number of atoms | 2376 | - | 2246 |
| Protein atoms | 2318 | - | 2219 |
| Ligand atoms | 12 | - | 27 |
| RMS bond lengths (Å) | 0.006 | - | 0.005 |
| RMS bond angles (°) | 0.82 | - | 0.96 |
| Ramachandran favored (%) | 98.0 | - | 91 |
| Ramachandran allowed (%) | 1.8 | - | 7.9 |
| Ramachandran outliers (%) | 0.35 | - | 0.72 |
| Clashscore | 12.09 | - | 14.04 |
| Mean *B* values (Å<sup>2</sup>) |  |  |  |
| Overall | 52.82 | - | 98.91 |
| Macromolecules | 52.87 | - | 99.09 |
| Ligands | 68.71 | - | 84.12 |
| Water | 46.16 | - | - |

<sup>a</sup>Parentheses numbers represent the highest-resolution shell.
<sup>b</sup> $R_{\text{merge}} = \sum_{hkl} \sum_i |I_i(hkl) - I_{av}(hkl)| / \sum_{hkl} \sum_i I_i(hkl)$ .

**Figure 2:**
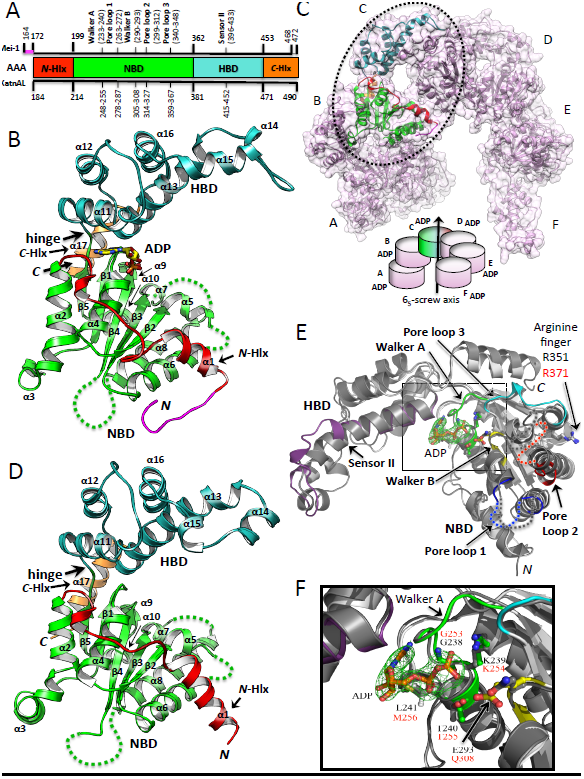
X-ray structures of ADP-bound MEI-1-AAA and monomeric nucleotide-free KATNAL1 (E308Q)-AAA. (A) Domain diagrams and numbering schemes for Katanin-AAA (colored according to the X-ray structures): N-terminal helix, N-Helix or N-Hlx; nucleotide binding domain, NBD; four-helix bundle domain, HBD; C-terminal helix, C-Helix or C-Hlx. Conserved motifs are shown in the schemes. (B) Ribbon diagram of MEI-1-AAA structure in the ADP state: N-terminal linker, magenta; N-Hlx, red; NBD, light green; HBD, cyan; C-Hlx, orange. The ADP is shown as stick representation. (C) The projection of MEI-1-AAA structure parallel to crystallographic 6_5_-screw axis shows a pseudo-hexameric left-handed spiral assembly. It is shown as ribbon diagram (light pink) with transparent surface representation background. One of the subunits is colored according to the crystal structure. (D) Ribbon representation of KATNAL1 (E308Q)-AAA in a nucleotide-free monomeric state. Domains are colored similar to MEI-1-AAA structure. (E) Overlay of MEI-1-AAA (pore loops 1-3, sensor II, walker A, walker B and backbone are colored in blue, red, cyan, purple, light green, yellow and dark gray, respectively) with the bound ADP molecule along its interacting residues shown as ball-and-stick representation, and KATNAL1 (E308Q)-AAA (white). The initial difference Fourier electron density contoured at 3 Σ for ADP is shown in green. (F) A close-up view of ADP binding pocket including the motifs shown in similar conformation as in panel E.

### Crystal structure of katanin’s AAA-ATPase in the apo-nucleotide state reveals a monomer

We determined the structure of human KATNAL1 (E308Q)/B1-con80 which grew in the space group *P*2_1_2_1_2_1_. We also observed degradation of MIT-con80 domains during crystallization similar to MEI-1/MEI-2. We determined the x-ray structure of KATNAL1 (E308Q) AAA ATPase using partial molecular replacement (MR) phases using the NBD domain of the MEI-1 AAA structure, combined with single anomalous dispersion (SAD) phases from a gold-derivative. The refined 2.4-Å x-ray structure reveals a single AAA-ATPase subunit in the asymmetric unit (Figure 2D; Figure S3C; Table 2), suggesting that the katanin AAA-ATPase is in a monomeric state, without nucleotide bound in the NBD pocket. In this structure, katanin does not form any typical AAA-ATPase oligomeric interfaces with neighboring AAA subunits (Figure S3C). The KATNAL1 (E308Q) AAA structure shows ordered density from residue 184 to residue 490 (α1-17 and β1-5) including the last four residues of C-terminal α17 (C-Hlx) which were absent from the MEI-1 AAA structure (Figure 2D; Figure S2). The structure shows elements in the nucleotide binding pocket including the Walker A and Walker B motifs in a similar conformation to those in the MEI-1 AAA structure, however, density for any nucleotide is completely absent in this structure (Figure 2E,F; Figure S2). The absence of any helical arrangement in the crystal is consistent with the SEC-MALS results indicating that KATNAL1 (E308Q) is a monomer in the absence of nucleotide (Figure 1D; Table 1). Thus this structure may represent the autoinhibited state of katanin.

### Structural comparison of katanin’s AAA-ATPase in the ADP and apo-nucleotide states reveal a conformational transition that inhibits oligomerization

We compared the MEI-1 AAA-ATPase structure in its pseudo-hexameric ADP state to the KATNAL1 (E308Q) AAA structure in its monomeric apo-nucleotide state through superimposing the two structures. The AAA-ATPase structures show a root-mean-square deviation (rmsd) of 1.50 Å (Cα positions) (Figure 3A,B). The comparison reveals the NBD fold becomes decompressed due to conformational changes in the N-helix, C-helix, and a refolding of the HBD domain. In the monomeric structure, the C-helix rotates 3°, while the second segment of the N-helix rotates 23° azimuthally (Figure 3A,B). The detailed interactions of N-helix and C-helix to NBD are shown in Figure S5. The HBD undergoes a significant refolding rearrangement in which sensor II elements are reorganized. In the ADP state, the central helix, α13, in the HBD is flanked by a parallel helix α15 near the tip of the HBD. However in the monomeric structure, the central helix in the HBD (α13) lengthens by two turns through refolding, while the parallel helix (originally α15) refolds into two two-turn helices (α15 and α14), which are oriented in nearly orthogonal orientations to α13. The HBD (α11-13 and α16) rotates 9° closer to the NBD. In this conformation, the HBD fold reorganizes and changes orientation leading to clockwise subdomain rotation (α15 and α14) (Figure 3A,B). The NBD in the monomeric structure shows decompression-like transitions compared to its fold in the pseudo-oligomeric structure, including a 12° rotation the α2 α3 helix-loop helix elements, and 10° α6 rotation. These NBD transitions involve elements mediating NBD-HBD interfaces in the pseudo-oligomeric state.

**Figure 3:**
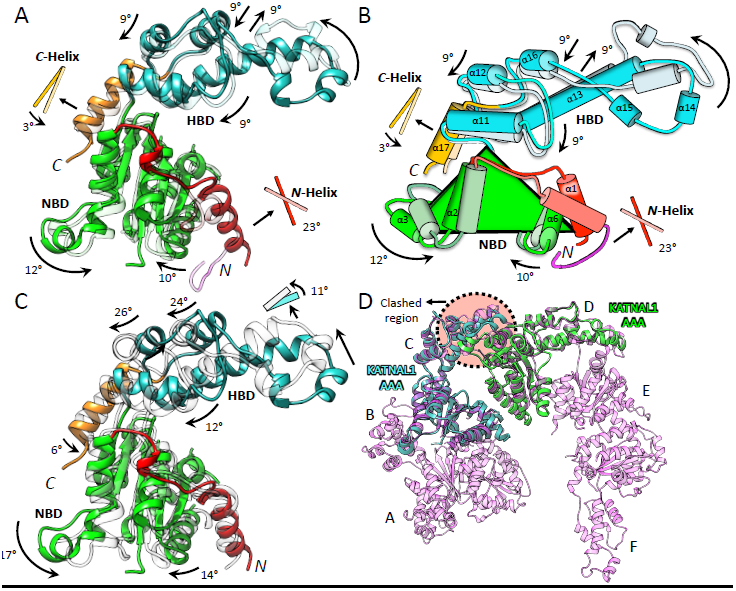
Structural comparison of Katanin-AAA subdomains and elements in monomeric and pseudo-oligomeric states. Arrows indicate the direction of rotational movements in all the panels. (A) Superimposition of KATNAL1 (E308Q)-AAA (color coded according to the structure) onto ADP-bound MEI-1-AAA (faded-colors) structure. (B) Schematic representation of the MEI-1-AAA (dark colors) overlaid on the KATNAL1 (E308Q)-AAA (faded-colors) as in panel A. (C) Superimposition of nucleotide-free state KATNAL1 (E308Q)-AAA (color-coded) onto ATP state KATNAL1-AAA structures. ATP model generated based on hexameric Vps4 structure (Monroe et al., 2017). (D) Superimposition of nucleotide-free KATNAL1 (E308Q) - AAA structures (cyan and green) onto two adjacent protomers in ADP-bound MEI-1 - AAA pseudo-hexameric left-handed spiral (light pink), with the backbones shown as ribbon diagrams. The clashed region (highlighted in circle with orange color) shows that the conformation of KATNAL1 (E308Q)-AAA is not compatible with pseudohexameric left-handed spiral assembly and it is in an auto-inhibited state.

To find out the difference between ATP and apo-nucleotide states, we built a homology model of KATNAL1 AAA-ATP state structure based on a Vps4 structure in an ATP-bound hexameric state (Monroe et al., 2017). We calculated those helical movements mentioned previously by superimposing the two structures (rmsd 2.36 Å; Cα positions), revealing that the rotational motions of these helices in the ATP state are amplified into slightly larger motions. The rotation angles are indicated in Figure 3C. These findings suggest that the conformation of katanin is strictly dependent on ATP or ADP binding. The overlay of the KATNAL1 (E308Q) AAA monomeric x-ray structure onto the MEI-1 AAA spiral assembly reveals that the HBD in the KATNAL1 (E308Q) AAA structure is not compatible with its docked state onto the NBD in the MEI-1 AAA structure. The conformation of the HBD in the monomeric state likely inhibits AAA pseudo-oligomeric assembly by interfering with its NBD interface (Figure 3D). The monomeric apo-nucleotide conformation is also incompatible with the hexameric ring assembly of Vps4 in the substrate-bound state (Monroe et al., 2017) (Figure S4D). The rmsd values between KATNAL1(E308Q) AAA and various katanin AAA ATPase ortholog structures are given in Table 3, revealing that ADP state MEI-1 AAA and apo-state KATNAL1(E308Q) AAA structures are in different conformations (Figure S4). The apo-nucleotide structure of KATNAL1 thus reveals the structural basis for autoinhibition.

**Table 3:**
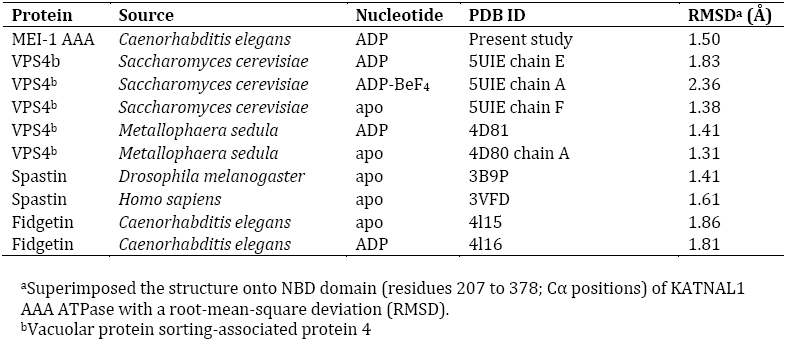
RMSD values of various AAA katanin orthologs.

| **Protein** | **Source** | **Nucleotide** | **PDB ID** | **RMSD<sup>a</sup> (Å)** |
| --- | --- | --- | --- | --- |
| MEI-1 AAA | *Caenorhabditis elegans* | ADP | Present study | 1.50 |
| VPS4<sup>b</sup> | *Saccharomyces cerevisiae* | ADP | 5UIE chain E | 1.83 |
| VPS4<sup>b</sup> | *Saccharomyces cerevisiae* | ADP-BeF<sub>4</sub> | 5UIE chain A | 2.36 |
| VPS4<sup>b</sup> | *Saccharomyces cerevisiae* | apo | 5UIE chain F | 1.38 |
| VPS4<sup>b</sup> | *Metallophaera sedula* | ADP | 4D81 | 1.41 |
| VPS4<sup>b</sup> | *Metallophaera sedula* | apo | 4D80 chain A | 1.31 |
| Spastin | *Drosophila melanogaster* | apo | 3B9P | 1.41 |
| Spastin | *Homo sapiens* | apo | 3VFD | 1.61 |
| Fidgetin | *Caenorhabditis elegans* | apo | 4I15 | 1.86 |
| Fidgetin | *Caenorhabditis elegans* | ADP | 4I16 | 1.81 |

<sup>a</sup>Superimposed the structure onto NBD domain (residues 207 to 378; Cα positions) of KATNAL1 AAA ATPase with a root-mean-square deviation (RMSD).
<sup>b</sup>Vacuolar protein sorting-associated protein 4

### Mapping *mei-1* mutations onto the full p60 katanin structural model

Using the sequenced *mei-1* loci (Clark-Maguire and Mains, 1994), we mapped the locations of these point mutations onto the full-length p60 structure, including both AAA-ATPase and MIT domains. The MIT domain of MEI-1 was modeled based on the recent crystal structure of *Mus musculus* p60N/p80C katanin complex (Jiang et al., 2017). The map of the *mei-1* mutations is shown in Figure 4 suggesting that the majority of the residues fall in the NBD. These mutations likely lead to its misfolding, or interference in its ability to bind ATP. Three of these *mei-1* mutations (sb3, sb23, ct103) lie at the junction of the NBD and HBD subdomains. One Mei1 mutation (ct89) lies in the HBD tip region, which we hypothesize is important for self-regulated monomer to oligomer conformational change. A single *mei-1* mutation (ct99) lies in the MIT domain and likely destabilize the MIT structure and interferes with MT binding. Only a single *mei-1* mutation (ct48) is exposed on the surface suggesting it may interfere with NBD-NBD assembly as formed in the katanin oligomers.

**Figure 4:**
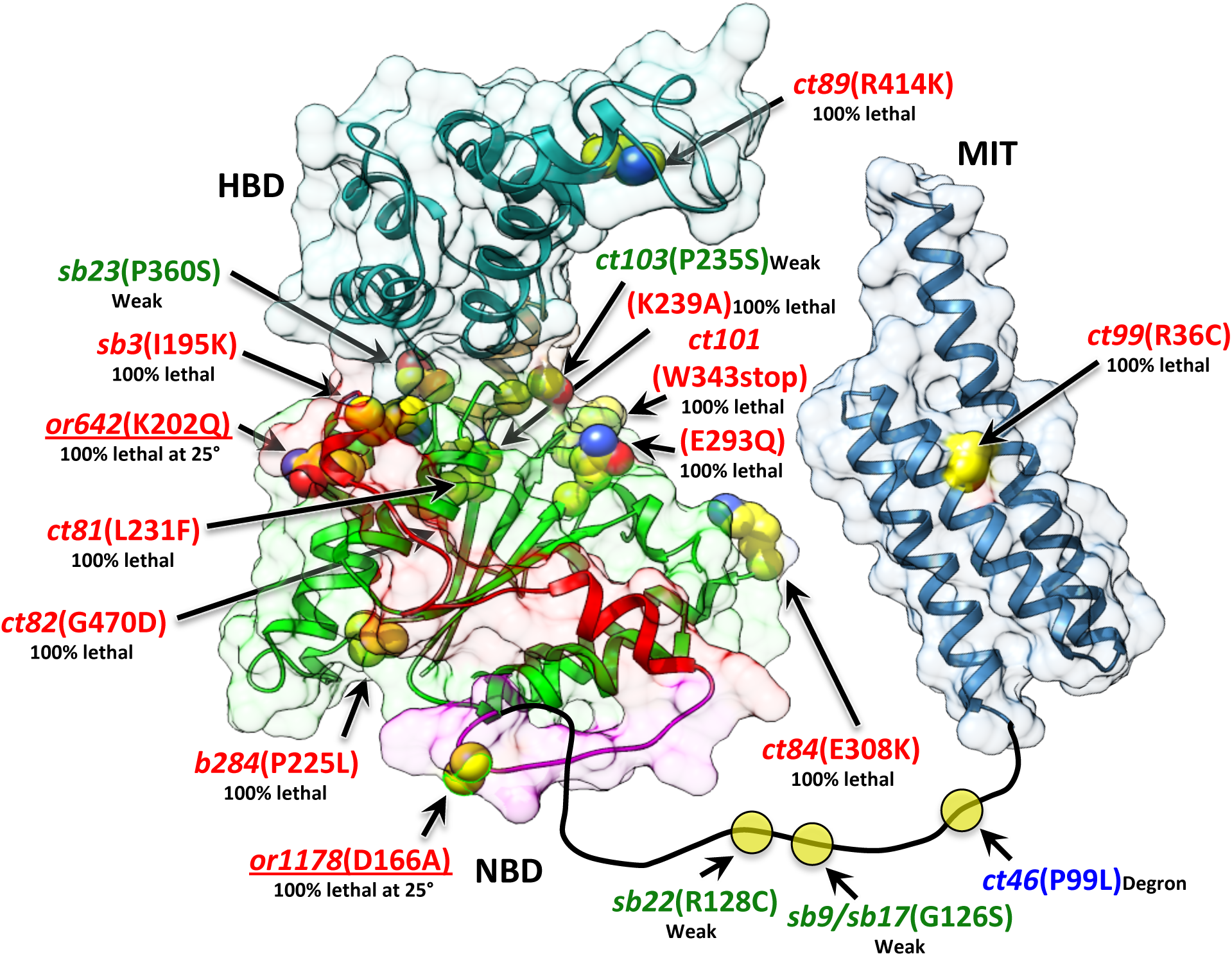
Mapping the *mei*1 *C. elegans* meiosis mutations onto structural full length MEI-1 P60 model: MEI-1-AAA (color coded according to the structure) and MIT (blue) are shown in ribbon representations with molecular transparent surface background and mutated residues are shown as yellow spheres with heteroatom colors. Phenotypes of functional mutations are labeled in different colors (100% lethal and no spindle poles, red; 100% lethal at 25°C, red with underlined; weak and partial loss of function, green; degron, blue). MIT domain was modeled based on *Mus musculus* MIT structure (Jiang et al., 2017).

### A model for katanin oligomerization and conformational changes involved in MT severing

We built a monomeric ATP-bound state of the AAA domain of human KATNAL1 using a VPS4 cryo-EM structure (Monroe et al., 2017) as a homology template and a monomeric ADP-bound state using the MEI-1 pseudo-hexameric structure as a homology template (Figure S6A,B). Using these monomeric ATP and ADP models, in addition to the autoinhibited apo-nucleotide X-ray structure, we generated three different hexameric structures by docking single subunits onto the Vps4 hexameric ring (Monroe et al., 2017) or the MEI-1 pseudo-hexameric left-handed spiral (Figure 2C). Docking all ATP-bound KATNAL1 protomers resulted in a hexameric right handed spiral (Figure S6C). This model terminates as a hexameric open ring because the small pitch of the helix does not allow addition of more protomers. This model is thus consistent with the discrete hexameric mass observed by SEC-MALS for KATNAL1(E308Q) in ATP (Figure 1E). Docking three adjacent ATP-bound protomers next to two ADP-bound protomers and an apo-nucleotide protomer resulted in a closed ring structure (Figure S6D) similar to that observed experimentally by cryo-EM for Vps4 (Monroe et al., 2017). The conformation of an apo-nucleotide KATNAL1 protomer reduces the size of the gap in the closed ring, thus preventing addition of a seventh protomer at either end. To explain why the pseudohexameric left-handed spiral model of ADP-KATNAL1 does not polymerize continuously in solution, we propose that one end could be capped by an apo-nucleotide protomer (Figure S6E). This apo-nucleotide capping leads to symmetry breaking. Difference between these hexameric conformations result specifically from variability in the HBD conformation (Figure S6D,E), which can contact with the NBD of a neighboring katanin leading to a closed hexamer, or remain in an open conformation depending on the nucleotide state of individual protomers. We suggest that sequential ATP hydrolysis and ADP release would propagate around the hexamer resulting in a cycle from right handed spiral, to closed ring, to left handed spiral and finally to dissociation of apo-nucleotide protomers as monomers.

## DISCUSSION

Using our two experimentally determined x-ray structures and three computationally built hexameric homology models (Figure S6D-F), we suggest a model for Katanin MT-based oligomerization and MT severing (Figure 5). We suggest that p60-p80 katanin heterodimers have a poor propensity for oligomerization due to the self-inhibited AAA conformation apparent in our KATNAL1 apo-nucleotide structure. Katanin heterodimers would bind along the MT lattice via the MIT-Con80 interface. These katanin p60-p80 heterodimers would diffuse along MT lattices, bind ATP and assemble into right handed open rings (Figure 5A-B). A right handed hexameric spiral of ATP-bound MEI-1 AAA domains was recently resolved by cryo-EM (Zehr et al., 2017). Previous fluorescence resonance energy transfer (FRET) experiments suggested that this monomer to oligomer transition requires both ATP binding and microtubule binding (Hartman and Vale, 1999). In vivo, assembly of right-handed open rings might be promoted by regulators such as ASPM (Jiang et al., 2017) or Patronin/CAMSAP (Jiang et al., 2014) which bind the MIT-con80 complex. Our monomeric KATNAL1 structure reveals the structural constraints that must be overcome to drive this initial assembly on the microtubule.

**Figure 5:**
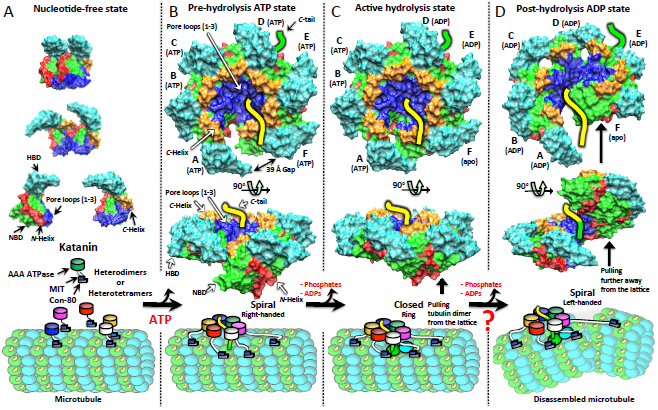
Structural model for Katanin AAA microtubule-based assembly and severing. (A) Upper panel: Nucleotide-free state Katanin AAA ATPase that form self-inhibited monomers or dimers in solution (Figure 2; N-Helix, red; NBD, light green; HBD, cyan; C-Helix, orange; pore loops1-3, blue). Lower panel: Cartoon diagrams of katanin p60-p80 (con80 domain) complexes diffusing on the MT lattice (α-tubulin, cyan; β-tubulin, light green; c-terminal tail or c-tail of β-subunit, yellow) in the absence of nucleotide. MIT (blue), Con80 (light purple) and AAA (green, magenta, blue, white and gold) domains are shown. (B) Upper panel: Side view of hexameric right-handed spiral model (produced based on Vps4, as described in figure S6) of Katanin AAA ATPase in pre-hydrolysis ATP state. Note that the pore loops are engaging the β-tubulin C-terminal tail (yellow) emanating from the MT lattice surface. Middle panel: Perpendicular view of the same assembly. Lower panel: Schematic diagram of heterododecameric p60-p80 right-handed spiral Katanin complex binding onto MT lattice via C-tail in the presence of ATP. (C) Upper panel: Side view of closed ring model (produced based on Vps4, as described in figure S6) of Katanin AAA ATPase in an active hydrolysis state. Middle panel: Perpendicular view of the same model. Lower panel: Ring structure of heterododecameric katanin complex displacing the tubulin dimer through c-tail from the MT lattice. (D) Upper panel: Side view of hexameric left-handed spiral model of Katanin AAA ATPase in post-hydrolysis ADP state (Figure 2). Middle panel: Perpendicular view of the same model. Lower panel: Cartoon representation of heterododecameric left-handed spiral Katanin complex pulling the tubulin dimer further away from the lattice leading to form two new MTs.

Katanin oligomers are thought to bind the β-tubulin C-termini through the AAA-ATPase pore loops located at the center of the hexamer (Bailey et al., 2015; Roll-Mecak and Vale, 2008). Initially, the open right-handed spiral of ATP-bound protomers would thread the β-tubulin C-terminal domain into its pore through the 39-Å gap in the ring. Upon hydrolysis of the ATP bound to three adjacent protomers and release of nucleotide from one protomer, the right-handed spiral would close into a flat ring encircling the beta tubulin tail (Figure 5C). This transition is similar to that proposed for VPS4 (Monroe et al., 2017; Su et al., 2017; Sun et al., 2017) and supported by a flat ring structure of MEI-1 determined by cryo-EM (Zehr et al., 2017). Upon hydrolysis of the ATPs associated with the remaining three protomers, the flat ring would transition to the left-handed spiral conformation (Figure 5D) with five ADP protomers and one apo-nucleotide protomer. In this state, NBD pore loops are repositioned further away from the microtubule leading to a pulling effect on the β-tubulin C-terminus away from the microtubule. This might induce the removal of an αβ-tubulin dimer from the MT lattice. Dissociation of ADP would result in a cycle back to monomers (Figure 5A). Removal of multiple tubulin dimers through multiple assembly cycles would generate a break in the microtubule. The proposed step-by-step changes in the AAA ATPase hexamers are shown in Figure S7B.

Zehr et al. (2017) suggested that the transition of a katanin hexamer from a right-handed open structure to a closed ring structure constitutes the power stroke for pulling a beta tubulin tail from the microtubule lattice. This transition, however, generates only a 15 Å displacement between pore loops. In contrast, the transition from a right handed helix to a left handed helix, as proposed in our model, would move pore loops over thrice this distance (∼50-Å). We suggest that this larger displacement is more likely to either denature the beta tubulin subunit or actually pull a tubulin dimer from the lattice.

The self-inhibition of katanin may be an important regulatory feature that prevents severing at inappropriate times and places at the low concentrations in the cytoplasm (estimated to be 20-50 nM). Alternatively, complete dissociation of hexamers into monomers might be an essential part of the microtubule-severing reaction. This is in contrast to other AAA-ATPases, such as Rho, NSF or p97, which are either stable hexamers or include a second AAA domain that stabilizes their oligomeric assembly (Adelman et al., 2006; Banerjee et al., 2016; Zhao et al., 2015). The structures presented here reveal the structural basis for this barrier to assembly.

## STRUCTURE DEPOSITION

Atomic coordinates and structure factors have been deposited in the Protein Data Bank with accession codes XXXX for KATNAL1 (E308Q) AAA and XXXX for MEI-1 AAA.

## AUTHOR CONTRIBUTIONS

S.N. did all of the experiments and contributed to experimental design and writing the manuscript. F.J.M. and J.A.B. contributed to experimental design and writing the manuscript.

## ACKNOWLEDGMENTS

We thank Advanced Photon Source (APS) and Dr. K. Rajashankar, J. Schuermann, and D. Neau of the Northeastern Collaborative Access Team (NE-CAT) in using the 24-IDC and IDE beam lines to collect all X-ray diffraction data for our crystallographic studies. We thank our graduate student, Mr. Brian D Cook for help with SEC-MALS. JAB and FJM are supported by National Institutes of Health GM110283 and GM079421, respectively.

## REFERENCES

Adelman, J.L., Jeong, Y.J., Liao, J.C., Patel, G., Kim, D.E., Oster, G., and Patel, S.S. (2006). Mechanochemistry of transcription termination factor Rho. Mol Cell 22, 611–621.

Ahmad, F.J., Yu, W., McNally, F.J., and Baas, P.W. (1999). An essential role for katanin in severing microtubules in the neuron. J Cell Biol 145, 305–315.

Babst, M., Wendland, B., Estepa, E.J., and Emr, S.D. (1998). The Vps4p AAA ATPase regulates membrane association of a Vps protein complex required for normal endosome function. EMBO J 17, 2982–2993.

Bailey, M.E., Sackett, D.L., and Ross, J.L. (2015). Katanin Severing and Binding Microtubules Are Inhibited by Tubulin Carboxy Tails. Biophys J 109, 2546–2561.

Banerjee, S., Bartesaghi, A., Merk, A., Rao, P., Bulfer, S.L., Yan, Y., Green, N., Mroczkowski, B., Neitz, R.J., Wipf, P., et al. (2016). 2.3 A resolution cryo-EM structure of human p97 and mechanism of allosteric inhibition. Science 351, 871–875.

Clark-Maguire, S., and Mains, P.E. (1994). Localization of the mei-1 Gene Product of Caenorhabditis elegans, a Meiotic-specific Spindle component. The Journal of Cell Biology 126, 11.

DeLaBarre, B., and Brunger, A.T. (2005). Nucleotide dependent motion and mechanism of action of p97/VCP. J Mol Biol 347, 437–452.

Fleming, K.G., Hohl, T.M., Yu, R.C., Muller, S.A., Wolpensinger, B., Engel, A., Engelhardt, H., Brunger, A.T., Sollner, T.H., and Hanson, P.I. (1998). A revised model for the oligomeric state of the N-ethylmaleimide-sensitive fusion protein, NSF. J Biol Chem 273, 15675–15681.

Fletcher, R.J., Bishop, B.E., Leon, R.P., Sclafani, R.A., Ogata, C.M., and Chen, X.S. (2003). The structure and function of MCM from archaeal M. Thermoautotrophicum. Nat Struct Biol 10, 160–167.

Hartman, J.J., and Vale, R.D. (1999). Microtubule Disassembly by ATP-Dependent Oligomerization of the AAA Enzyme Katanin. Science 286, 4.

Hazan, J., Fonknechten, N., Mavel, D., Paternotte, C., Samson, D., Artiguenave, F., Davoine, C.S., Cruaud, C., Durr, A., Wincker, P., et al. (1999). Spastin, a new AAA protein, is altered in the most frequent form of autosomal dominant spastic paraplegia. Nat Genet 23, 296–303.

Hu, W.F., Pomp, O., Ben-Omran, T., Kodani, A., Henke, K., Mochida, G.H., Yu, T.W., Woodworth, M.B., Bonnard, C., Raj, G.S., et al. (2014). Katanin p80 regulates human cortical development by limiting centriole and cilia number. Neuron 84, 1240–1257.

Iwaya, N., Kuwahara, Y., Fujiwara, Y., Goda, N., Tenno, T., Akiyama, K., Mase, S., Tochio, H., Ikegami, T., Shirakawa, M., et al. (2010). A common substrate recognition mode conserved between katanin p60 and VPS4 governs microtubule severing and membrane skeleton reorganization. J Biol Chem 285, 16822–16829.

Jiang, K., Hua, S., Mohan, R., Grigoriev, I., Yau, K.W., Liu, Q., Katrukha, E.A., Altelaar, A.F., Heck, A.J., Hoogenraad, C.C., et al. (2014). Microtubule minus-end stabilization by polymerization-driven CAMSAP deposition. Dev Cell 28, 295–309.

Jiang, K., Rezabkova, L., Hua, S., Liu, Q., Capitani, G., Altelaar, A.F.M., Heck, A.J.R., Kammerer, R.A., Steinmetz, M.O., and Akhmanova, A. (2017). Microtubule minus-end regulation at spindle poles by an ASPM-katanin complex. Nat Cell Biol 19, 480–492.

McNally, F.J., and Thomas, S. (1998). Katanin is responsible for the M-phase microtubule-severing activity in Xenopus eggs. Mol Biol Cell 9, 1847–1861.

McNally, F.J., and Vale, R.D. (1993). Identification of katanin, an ATPase that severs and disassembles stable microtubules. Cell 75, 419–429.

McNally, K., Audhya, A., Oegema, K., and McNally, F.J. (2006). Katanin controls mitotic and meiotic spindle length. J Cell Biol 175, 881–891.

Mishra-Gorur, K., Caglayan, A.O., Schaffer, A.E., Chabu, C., Henegariu, O., Vonhoff, F., Akgumus, G.T., Nishimura, S., Han, W., Tu, S., et al. (2014). Mutations in KATNB1 cause complex cerebral malformations by disrupting asymmetrically dividing neural progenitors. Neuron 84, 1226–1239.

Monroe, N., Han, H., Shen, P.S., Sundquist, W.I., and Hill, C.P. (2017). Structural basis of protein translocation by the Vps4-Vta1 AAA ATPase. eLife 6.

Roll-Mecak, A., and McNally, F.J. (2010). Microtubule-severing enzymes. Curr Opin Cell Biol 22, 96–103.

Roll-Mecak, A., and Vale, R.D. (2008). Structural basis of microtubule severing by the hereditary spastic paraplegia protein spastin. Nature 451, 363–367.

Scott, A., Chung, H.Y., Gonciarz-Swiatek, M., Hill, G.C., Whitby, F.G., Gaspar, J., Holton, J.M., Viswanathan, R., Ghaffarian, S., Hill, C.P., et al. (2005). Structural and mechanistic studies of VPS4 proteins. EMBO J 24, 3658–3669.

Sharma, N., Bryant, J., Wloga, D., Donaldson, R., Davis, R.C., Jerka-Dziadosz, M., and Gaertig, J. (2007). Katanin regulates dynamics of microtubules and biogenesis of motile cilia. J Cell Biol 178, 1065–1079.

Srayko, M., Buster, D.W., Bazirgan, O.A., McNally, F.J., and Mains, P.E. (2000). MEI- 1/MEI-2 katanin-like microtubule severing activity is required for Caenorhabditis elegans meiosis. Genes Dev 14, 1072–1084.

Su, M., Guo, E.Z., Ding, X., Li, Y., Tarrasch, J.T., Brooks, C.L., 3rd, Xu Z., and Skiniotis, G. (2017). Mechanism of Vps4 hexamer function revealed by cryo-EM. Sci Adv 3, e1700325.

Sun, S., Li, L., Yang, F., Wang, X., Fan, F., Yang, M., Chen, C., Li, X., Wang, H.W., and Sui, S.F. (2017). Cryo-EM structures of the ATP-bound Vps4E233Q hexamer and its complex with Vta1 at near-atomic resolution. Nat Commun 8, 16064.

Suno, R., Niwa, H., Tsuchiya, D., Zhang, X., Yoshida, M., and Morikawa, K. (2006). Structure of the whole cytosolic region of ATP-dependent protease FtsH. Mol Cell 22, 575–585.

Wohlever, M.L., Mateja, A., McGilvray, P.T., Day, K.J., and Keenan, R.J. (2017). Msp1 Is a Membrane Protein Dislocase for Tail-Anchored Proteins. Mol Cell 67, 194–202 e196.

Zehr, E., Szyk, A., Piszczek, G., Szczesna, E., Zuo, X., and Roll-Mecak, A. (2017). Katanin spiral and ring structures shed light on power stroke for microtubule severing. Nat Struct Mol Biol.

Zhao, M., Wu, S., Zhou, Q., Vivona, S., Cipriano, D.J., Cheng, Y., and Brunger, A.T. (2015). Mechanistic insights into the recycling machine of the SNARE complex. Nature 518, 61–67.

